# Characterization of a hypersporulating strain derivative of *Clostridioides difficile* R20291

**DOI:** 10.1101/2025.11.25.690273

**Authors:** Francisca Cid-Rojas, Daniel Paredes-Sabja

**Author notes:** Corresponding Author. Dr. Daniel Paredes-Sabja, Department of Biology, Texas A&M University, College Station, TX, 77843, USA.

## Abstract

*Clostridioides difficile* is a Gram-positive and obligate anaerobic spore-former pathogen. *C. difficile* spores are essential for the transmission and recurrence of *C. difficile* infections (CDI). A major challenge in sporulation studies in *C. difficile* is the low yield and asynchronous nature of this process. In this work, a hypersporulating strain, derivative of R20291, with an early sporulation onset and enhanced sporulation efficiency was isolated by serendipity. This strain had 1000-fold higher sporulation efficiency than the parental R20291 strain in sessile culture conditions. Electron micrographs revealed that spores of both strains have similar hair-like projections, electron-dense outer exosporium layer features. Whole genome sequencing and genomic analyses revealed that the hypersporulating strain had a 2356 bp-deletion spanning three ORF, including a non-essential *proC1* involved in proline metabolism, and a missense mutation in *rsbV*, an anti-anti-SigB factor of RsbW. These observations suggest that this RsbV-variant might contribute to constitutive repression of the SigB-dependent general stress response, and therefore, derepressing sporulation in this hypersporulating strain.

## Introduction

*Clostridioides difficile* is an anaerobic, Gram-positive, spore-forming pathogen and the principal etiologic agent of *C. difficile* infection (CDI), as well as, currently being the most frequent healthcare-associated cause of infectious diarrhea in industrialized countries (1, 2). CDI typically occurs after antibiotic-mediated disruption of the normal gut microbiota, enabling endogenous or ingested *C. difficile* spores to germinate and colonize (3, 4). Clinical manifestations range from mild diarrhea to pseudomembranous colitis, toxic megacolon and, in severe cases, death (3, 4). Moreover, recurrent CDI (R-CDI) remains a major public-health challenge, which affects 20-30% of CDI patients and may increase to nearly 60% after multiple episodes (5, 6).

The infection cycle of *C. difficile* is primarily driven by its ability to produce metabolically dormant and environmentally resilient spores (7). After ingestion, spores survive the passage through the stomach and germinate in the gastrointestinal tract in response to bile acids and other cues (7, 8), leading to colonization, toxin production, and disease development (9). A subset of cells initiate sporulation, generating new spores that can persist in the host and are shed into the environment (10).

Although the core regulatory stages of sporulation have been characterized in *C. difficile*, sporulation-related studies are challenging due to low sporulation frequency under laboratory conditions (11-14), and that cultures undergo sporulation in an inherently asynchronous manner (12, 15). Unlike other genetically trackable spore-formers, such as *B. subtilis* and *C. perfringens*, the heterogeneity and low sporulation frequency in *C. difficile*, complicates detailed investigations of sporulation-specific phenotypes, limits the ability to obtain stage-enriched populations, and reduces yields of purified spores for biochemical, structural, or high-throughput assays. Attempts to create high sporulating strains in *C. difficile* have met partial success, such as overexpression of Spo0A (11), however, tunning sporulation commitment requires consideration of additional regulators such as the phosphatase Spo0E and RtsA, both known to act as repressors by sequestering Spo0A (16, 17).

In the context of unrelated culturing of *C. difficile*, we serendipitously identified a derivative of the epidemic lineage strain R20291 that exhibits a distinguishable hypersporulation phenotype. In this study, we characterize this hypersporulating strain (R20291_CM196_) and assess its potential as a laboratory tool for generating high-yield and more synchronous spore preparations. We observed that R20291_CM196_ presented a 30 to 1,000-fold increase in ethanol-resistant CFU, depending on the sporulation media composition, and reached a total of ∼ 60 % sporulating cells after 24 h of incubation in 70:30 agar plates. Moreover, electron micrographs show that spores from the hypersporulating strain are identical to parental strain. Whole genome sequence revealed the presence of a ∼2000 bp deletion including a pyrroline-carboxylate reductase, *proC1*, involved in proline biosynthesis and a single nucleotide polymorphism in *rsbV*, an anti-anti-SigB factor, involved in the general stress response and inhibition of sporulation by Spo0E activation. Because this sporulation phenotype occurs under *in vitro* conditions, the strain offers a practical platform for detailed studies of *C. difficile* sporulation and spore physiology, where higher frequency of individual developmental stages is required.

## Materials and Methods

### Bacterial strains and growth conditions

*C. difficile* strains were routinely grown at 37□°C under anaerobic conditions in a Coy Vinyl Anaerobic chamber (COY laboratory products, USA) in BHIS medium: 3.7% weight□vol^−1^ brain heart infusion (BD, USA) supplemented with 0.5% weight□vol^−1^ yeast extract (BD, USA) and 0.1% weight□vol^−1^ L-cysteine (Thermo Fisher Scientific, USA); TY medium: 3% weight□vol^−1^ trypticase (BD, USA) with 0.5% weight□vol^−1^ yeast extract (BD, USA) or 70:30 medium: 6.3% weight□vol^−1^ peptone (BD, USA), 0.35% weight□vol^−1^ protease peptone (BD, USA), 0.07% weight□vol^−1^ ammonium sulfate (NH_4_)_2_SO_4_ (Merck USA), 0.106% weight□vol^−1^ Tris base (Omnipur, Germany), 1.11% weight□vol^−1^ brain heart infusion (BD, USA) and 0.15% weight□vol^−1^ yeast extract (BD, USA). To grow strains in plates, 1.5% weight□vol^−1^ agar (VWR, USA) was added to medium. Strains used in this work are summarized in Table S1.

### Growth assays

*C. difficile* strains were cultured anaerobically on pre-reduced BHIS broth supplemented with 0.1% sodium taurocholate (NaTC, Sigma, USA) and 0.5% glucose (BHISG + NaTC) for 16 h at 37 ºC to prevent sporulation. Overnight culture was diluted 1:3 with pre-warmed BHISG + NaTC broth and incubated for 2 h at 37 ºC to allow cells to enter exponential phase. Subsequently, culture was diluted 1:100 in fresh BHIS and 100 μL were added to wells in a 96-well plate. Optical density (OD_600_) was measured every 3 minutes for 22 hours at 37 ºC using Cerillo’s Alto™ microplate reader. Three independent biological replicates with four technical replicates each, were performed per sample.

### Biofilm formation

Biofilm formation was measured as previously described (18, 19). Briefly, *C. difficile* inoculated BHISG broth was diluted 1:50, 1□mL aliquots were added to 24-well plate and incubated at 37□°C in anaerobic conditions for 24□h. Biofilm biomass was stained with 0.1% crystal violet (CV) for 1h, gently washed with 1X PBS and extracted with 100% ethanol for 2 h at room temperature. Solubilized CV was diluted 1:10 and absorbance was measured at OD_570_ with plate reader (BioTek, Synergy H1).

### Spore purification

As previously described (19, 20), 250 µL of a *C. difficile* inoculated BHISF (0.2% Fructose) + NaTC broth was plated on pre-reduced 70:30 agar plates (35 mL) and incubated for 5 days at 37□°C in a COY anaerobic chamber. Sporulating culture was harvested and cleaned with ice-cold sterile Milli-Q water. Nycodenz (Axell USA) 45% solution was used to purified spores by centrifugation at 18,400 × g for 20 min, cellular debris was removed, spore pellet separated and washed with sterile Milli-Q water and stored at −80□°C until use.

### Whole genome sequence

*C. difficile* strains were whole genome sequenced (SeqCenter, USA) and Illumina data was analyzed using Geneious Prime^®^ 2025.1.3. Reads were mapped to *C. difficile* R20291 reference genome (GenBank: CP029423) using Bowtie2 v7.2.2, with default parameters. Whole genome variant calling was performed by Geneious variant finder with a minimum coverage of five reads and a minimum variant frequency of 0.95 and long INDEL (insertion or deletion) regions were predicted using Find Low/High Coverage tool.

### Sporulation assay by ethanol resistance and phase-contrast microscopy

*C. difficile* sporulation efficiency was assayed as previously described (19, 21). Briefly, plating was performed as previously described in spore purification methods in BHIS, TY and 70:30 agar plates. Cells were harvested after 24 h in 1X PBS, OD_600_ was measured and absorbance of sporulating culture was adjusted to 1. An aliquot of 250 μL was treated with 100% ethanol for 15 minutes at room temperature, serially diluted and spot plated in triplicate onto BHIS + NaTC plates under anaerobic conditions. Sporulation frequency was calculated as the proportion of spores that germinated after ethanol treatment, divided by the total number of CFU in the untreated sample. The ratio of vegetative cells vs spores/sporulating cells was also determined using phase contrast microscopy (Leica DMRX; Germany).

For sporulation timing, cultures were plated in 70:30 plates and incubated at 37 ºC under anaerobic conditions. Samples were harvested after 8, 10, 12, 14, 16, 18, 20 and 24 h into 1 mL 1X PBS. Cells were centrifuged at 18,400□×□g for 5□min and fixed with 4% paraformaldehyde. Vegetative and spores/sporulating cells count was determined using phase contrast microscopy (Leica DMRX; Germany).

### Transmission electron microscopy (TEM)

As previously described with some modifications (10, 22), spores were fixed overnight with 2.5% glutaraldehyde and 1% paraformaldehyde in 0.1□M cacodylate buffer. Secondary fixation was performed with 1% osmium tetroxide for 2 h, followed by 30 min staining with 1% tannic acid. Samples were serially dehydrated 30 min each with acetone 30% (with 2% uranyl acetate), 50%, 70%, 90%, and twice with 100% acetone, after which samples were embedded in Spurr resin (22, 23). Thin sections (90 nm) were obtained using a microtome, placed on glow-discharge carbon-coated grids, and double-stained with 2% uranyl acetate followed by lead citrate. Spores were analyzed using a Philips Tecnai 12 Bio Twin microscope at the Unidad de Microscopía Avanzada, Pontificia Universidad Católica de Chile.

### Statistical analyses

Statistical significance was determined by Welch’s t-test in GraphPad Prism v9.0. <0.05 (*), <0.01 (**), <0.001 (***), <0.0001 (****), ns: no significance. All experiments were performed in triplicate with at least three technical replicates.

## Results and discussions

### Hyper-sporulating R20291 derivative

A canonical feature of *C. difficile* spore formation is its asynchronous nature and low yield. By serendipity, in a routine experiment that an isolate of *C. difficile* R20291 (from here on address as R20291_CM196_) presented a marked increase in the number of spores after 24 h incubation in 70:30 sporulation media in comparison with the usual levels obtained in lab with this strain (wild-type strain from here on address as R20291_CM210_) (**Fig. 1A**).

**Fig. 1.**
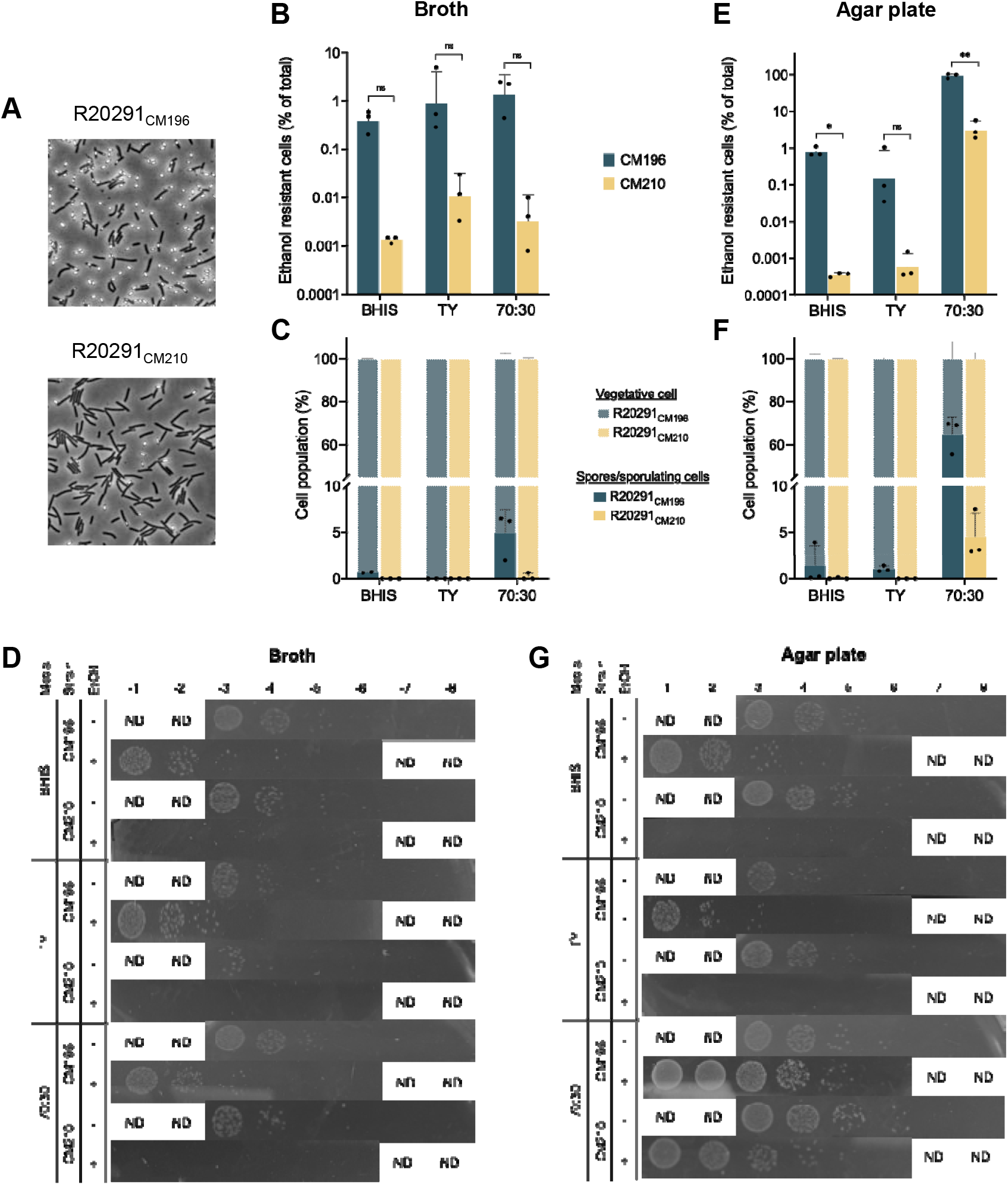
Hypersporulating *C. difficile* R20291 derivative strain. (A) Phase contrast microscopy of a 24 h sporulating culture. Sporulation efficiency evaluated by ethanol resistant sporulation assay of 24 h sporulating cultures grown in BHIS, TY or 70:30 media (B) broth or (E) agar. Sporulation efficiency evaluated by phase contrast microscopy images of 24 h sporulating cultures grown in BHIS, TY or 70:30 media (C) broth or (F) agar. At least 800 spores/cells were counted from each sample. Representative images of serial spot dilution of ethanol resistant sporulation assay in (D) broth and (G) agar plate. Cells were spotted at 10-fold serial dilutions on BHIS agar plates supplemented with 0.1% sodium taurocholate. ND: no data. Welch’s t-test. < 0.05 (*), < 0.01 (**), ns: no significance.

To confirm and evaluate if the increased amounts of spores observed in R20291_CM196_ strain were confined to specific nutritional conditions, a sporulation efficiency assay by ethanol resistance of cells grown in three different media (BHIS, TY and 70:30) widely used in *C. difficile* research was conducted. There was no significant difference in the ethanol-resistant CFU by these two strains when grown in any broth media (*p* > 0.05) (**Fig. 1B,1D and S1**). However, when cells were plated in 70:30 and BHIS agar plates, but no in TY agar plates, a significant difference (p < 0.01 and p < 0.05, respectively) in ethanol-resistant CFU was observed. This difference was highest in 70:30 agar plates, sporulation efficiency where ∼ 92% for the hypersporulating R20291_CM196_ and 3% for the parental R20291_CM210_ strain, respectively (**Fig. 1E, 1G and S2**).

This phenotype was further assessed by phase-contrast microscopy. In 70:30 broth, strain R20291_CM196_ produced 4.9% phase-bright spores, whereas growth on 70:30 agar resulted in a marked increase to 65%. By comparison, R20291_CM210_ displayed substantially lower sporulation frequencies, with 0.2% spores in broth and 4.5% on agar (**Fig. 1C and 1F**). The higher proportion of ethanol-resistant CFU relative to microscopic counts observed for R20291_CM196_ (92% vs. 65%) likely reflects loss of viability of vegetative cells during sporulation, resulting in their underrepresentation in ethanol-resistance assays but continued detection by microscopy.

### R20291_CM196_ exhibits an early onset of sporulation

To determine whether R20291_CM196_ and R20291_CM210_ initiated sporulation at similar times, sporulation-timing assay. Cultures grown on 70:30 agar were harvested at defined intervals was performed and examined by phase-contrast microscopy. Free spores were first detectable in R20291_CM196_ at 12 h, approximately 2 h earlier than in R20291_CM210_, and were ∼7-fold more abundant at this time point (7% vs. 1%, respectively) (**Fig. 2A and 2B**). By 14 h, R20291_CM196_ reached 31% spores compared to 1% in R20291_CM210_. After 24 h, more than half of the R20291_CM196_ population consisted of phase-bright spores (57%), representing a ∼10-fold higher sporulation level than observed in R20291_CM210_ (5%). These results indicate that R20291_CM196_ not only initiates sporulation earlier but that a larger fraction of the population commits to sporulation over time compared with R20291_CM210_. Because individual cell-transition rates cannot be directly measured under these conditions, the precise rate of sporulation remains undetermined.

**Fig. 2.**
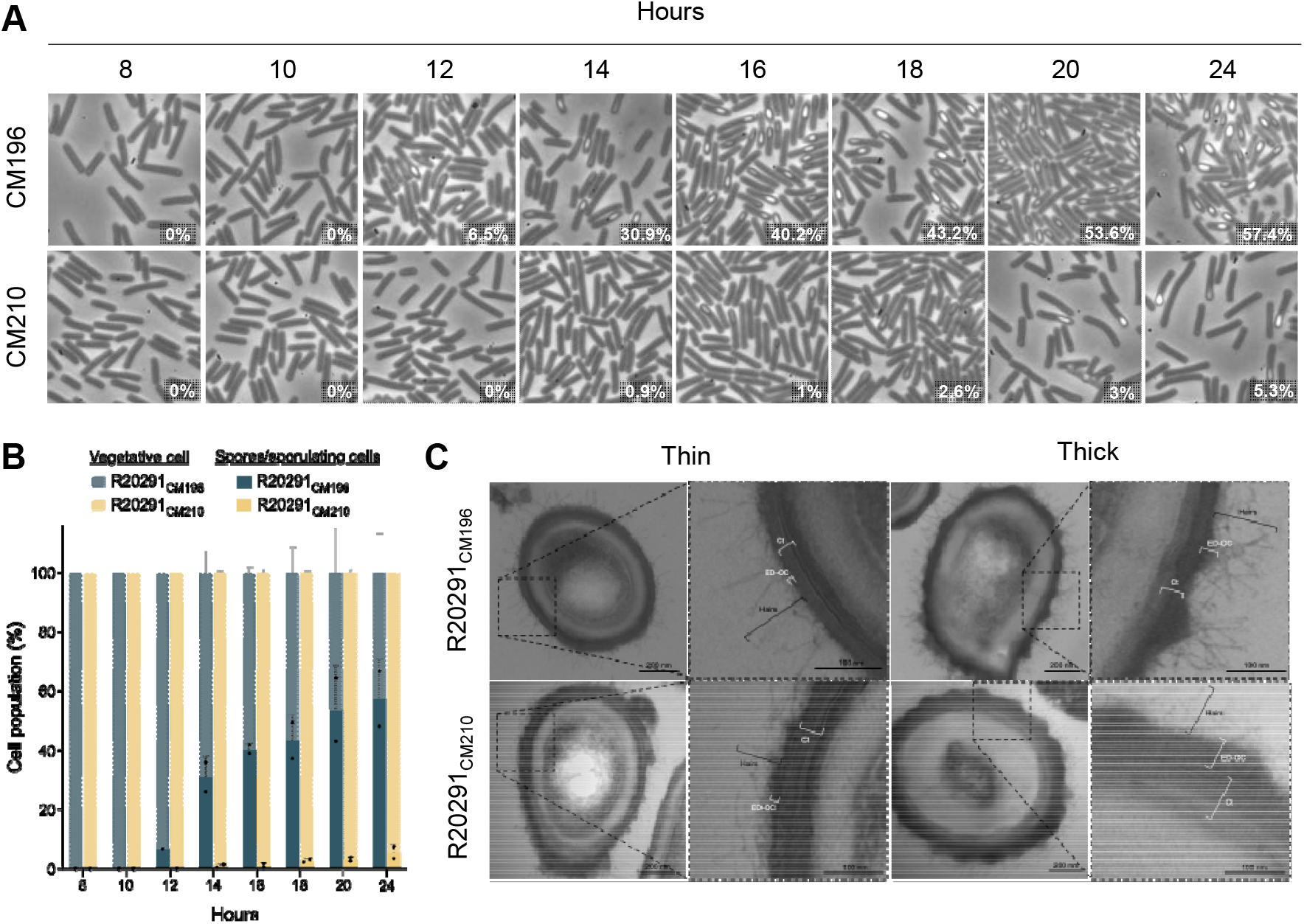
Early sporulation onset and spore ultrastructure analysis of R20291_CM196_. (**A**) Sporulation timing in 70:30 agar plates. Sporulating culture was monitored at 8, 10, 12, 14, 16, 18, 20 and 24 h. (**B**) Representative phase contrast microscopy images of sporulation timing assay. Numbers on the lower right corner corresponds to the percentage of spores/sporulating cells in the sample at that time. (**C**) Transmission electron microscopy (TEM) of R20291_CM196_ and R20291_CM210_ spores. ED-OC: Electron-dense outer coat. Ct: Coat.

### Ultrastructural analysis of R20291_CM196_ spores

To assess whether the hypersporulation phenotype altered spore ultrastructure, transmission electron microscopy was performed. Spores produced by R20291_CM196_ displayed no detectable structural abnormalities and retained characteristic morphological features, including hair-like projections over the electron-dense outer coat (ED-OC) and the lamellar organization of the coat (Ct) in both thin- and thick-coat morphotypes (**Fig. 2C**). These observations indicate that the hypersporulation phenotype of R20291_CM196_ affects the timing and frequency of sporulation but does not appear to impact spore ultrastructure.

### R20291_CM196_ exhibit differences in growth and biofilm formation

To further characterize the physiology of R20291_CM196_, planktonic growth and biofilm formation was examined. R20291_CM196_ exhibited reduced growth, reaching a maximal OD_600_ of 0.1 compared with 0.2 for R20291_CM210_. The onset of the decline phase also occurred earlier in R20291_CM196_, beginning at approximately 10 h post-inoculation, whereas R20291_CM210_ entered decline around 18 h (Fig. 3A). Biofilm formation was significantly diminished in R20291_CM196_, which produced an OD_570_ of 0.3 compared with 0.6 for R20291_CM210_ (p < 0.0001) (Fig. 3B). These data suggest that the increased commitment to sporulation observed in R20291_CM196_ may influence both planktonic growth dynamics and the capacity for form sessile communities.

**Fig. 3.**
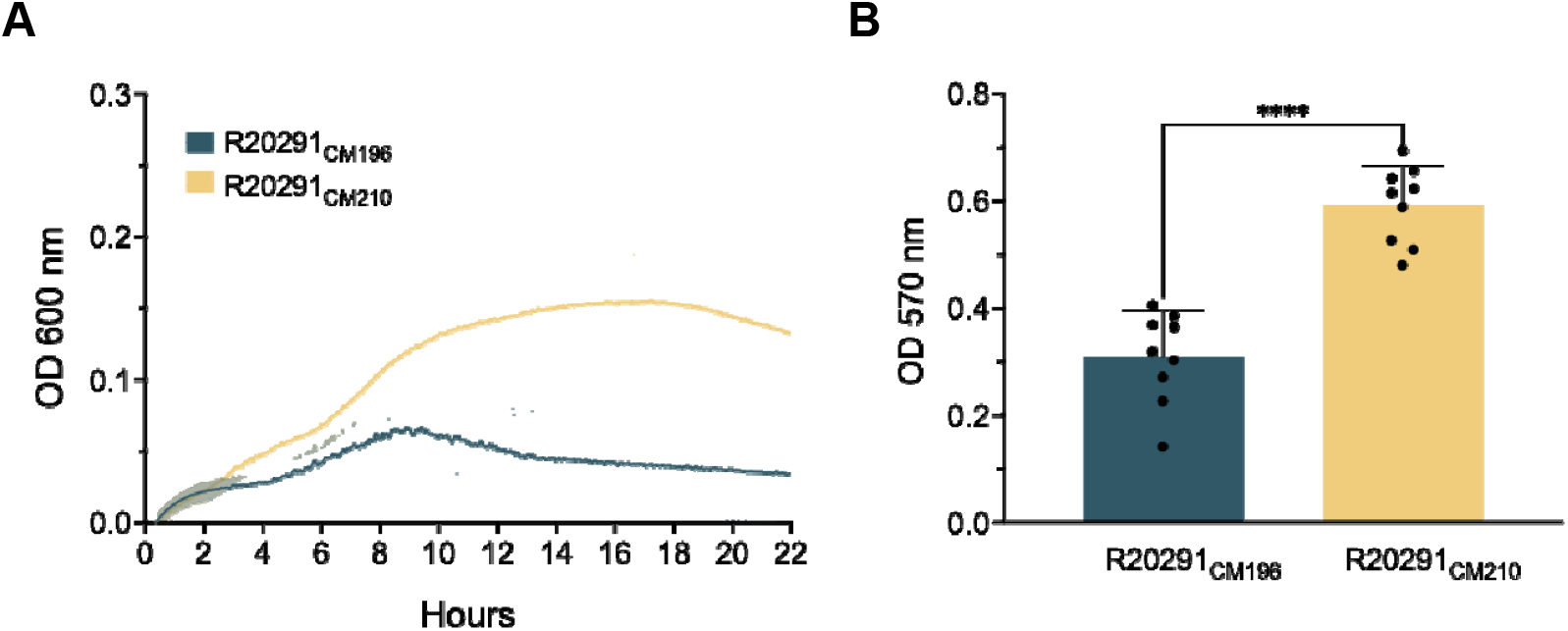
R20291_CM196_ physiology traits. (**A**) Growth curve of R20291_CM196_ (teal) and R20291_CM210_ (yellow) in BHIS media over 22 h. (**B**) Biofilm formation was evaluated at 24 h by crystal violet staining and measured at OD_570_. Welch’s t-test. < 0.0001 (****)

### Whole genome sequencing of R20291_CM196_ reveals unique polymorphism

To determine whether the differences in sporulation commitment in R20291_CM196_ were attributed to underlying genetic variation, R20291_CM196_ and R20291_CM210_ were whole genome sequenced and subjected to comparative genomics against *C. difficile* reference genome CP029423. Both isolates shared five mutations relative to the reference strain. These included two missense mutations: one in *rsbW* (G82V, CDIF27147_00009), which encodes the anti-sigma factor serine protease of SigB, and another in *vncR* (D202G, CDIF27147_02036), a response regulator of a two-component system (Fig. 4A). In addition, two nonsense mutations resulting from single-nucleotide deletions were identified in *rbsK* (CDIF27147_00429), encoding ribokinase, and in a two-component sensor histidine kinase (CDIF27147_02733) (Fig. 4A); Finally, a single nucleotide insertion in a non-coding intergenic region located 336 bp downstream of CDIF27147_00700 and 144 bp upstream of CDIF27147_00701 was detected (**Fig. 4A**).

**Fig. 4.**
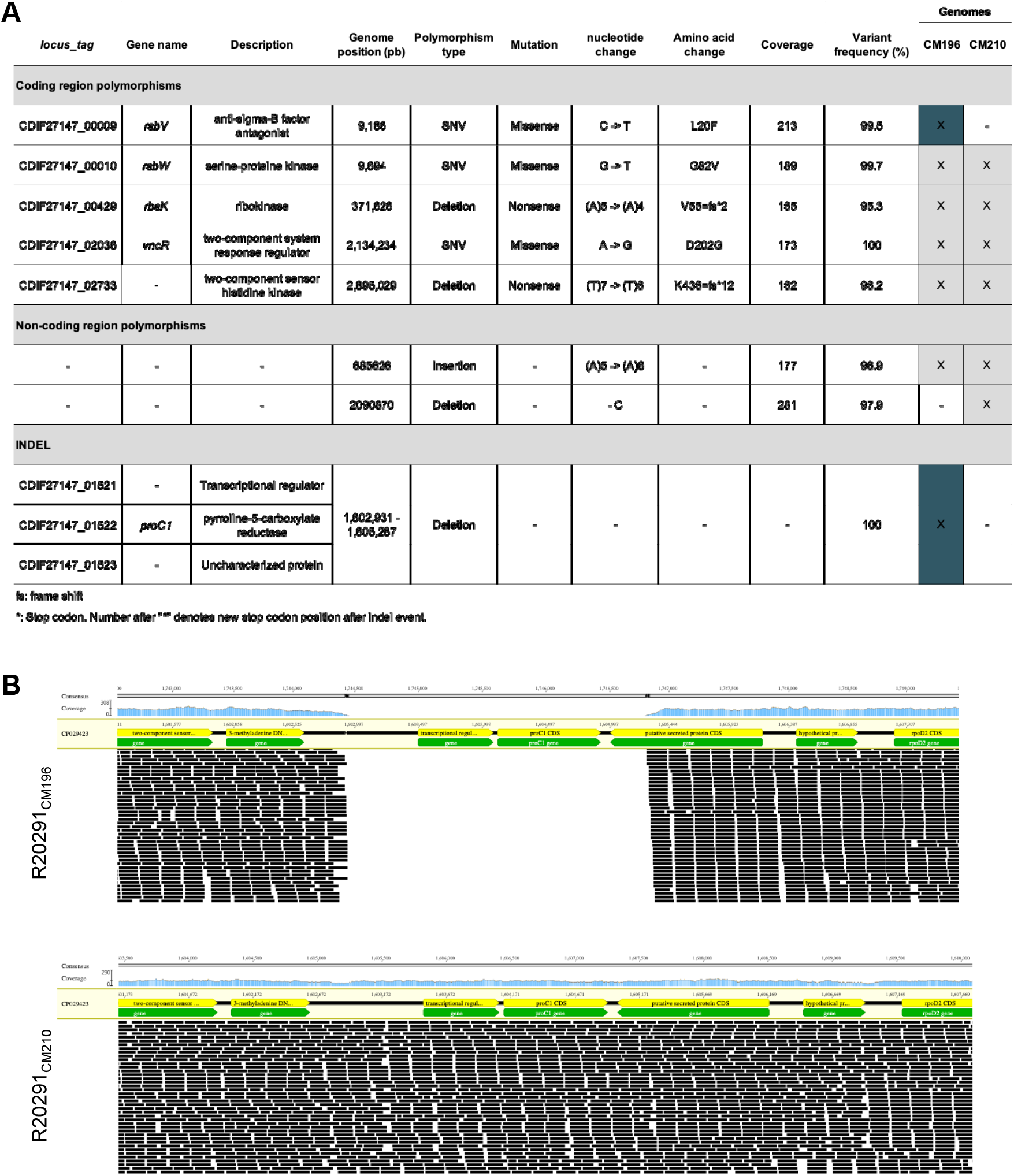
Mutations in hypersporulating *C. difficile* R20291_CM196_ strain. (**A**) Description of single nucleotide variants and INDELs present in R20291_CM196_ and R20291_CM210_ strains in comparison with *C. difficile* R20291 reference genome (CP029423). (**B**) Genetic deletion in R20291_CM196_ vs wildtype region in R20291_CM210_.

Notably, R20291_CM196_ harbors two additional polymorphisms not present in R20291_CM210_. One of these is a missense mutation (L20F) in *rsbV*, which encodes the anti-anti-sigma factor that regulates SigB and plays a crucial role in the general stress response against pH stress, antimicrobial peptides and reactive oxygen species (ROS) (24, 25). In *B. subtilis*, RsbV functions by binding and sequestering RsbW, thereby releasing SigB from the RsbW-SigB complex and allowing for the activation of the SigB regulon (26). Activation of SigB has been shown to reduce sporulation in *B. subtilis* (27), in part through upregulation of *spo0E*, which encodes an aspartyl phosphatase that dephosphorylates the master of sporulation Spo0A (27, 28). In *C. difficile spo0E* mutants exhibit twofold increase in sporulation relative to wild-type, consistent with Spo0E-mediated repression of sporulation observed in *B. subtilis* (16). Because sporulation is an energetically costly process and generally triggered only after other stress-response pathways have been exhausted (29), cells typically activate the general stress response prior to committing to sporulation. The L20F substitution in RsbV may impair its ability to bind RsbW, thereby allowing RsbW to remain associated with SigB. Under this scenario, SigB would remain inhibited, limiting activation of the SigB regulon and its downstream repression of sporulation.

The second mutation unique to R20291_CM196_ is a 2,356 bp deletion encompassing three adjacent genes: a transcriptional regulator (CDIF27147_01521), a pyrroline-5-carboxylate reductase (*proC1*, CDIF27147_01522) involved in the final step of L-proline biosynthesis from glutamate, and the C-terminal 284 bp of an uncharacterized protein (CDIF27147_01523) (**Fig. 4A and 4B**). In *B. subtilis*, only the simultaneous loss of the three pyrroline-5-carboxylate reductases (*proG, proH*, and *proI*) results in proline auxotrophy, whereas the individual genes are not essential (30). The *C. difficile* R20291 genome encodes two predicted pyrroline-5-carboxylate reductases, *proC1* and *proC2*(CDIF27147_03465). These proteins share 29–33% identity with *B. subtilis* ProH/ProI and approximately 23% identity with ProG. Thus, deletion of *proC1* alone is unlikely to abolish proline biosynthesis in R20291CM196, as *proC2* remains intact. The deletion also removes CDIF27147_01521, the putative transcriptional regulator, and truncates CDIF27147_01523; however, the functions of these genes remain unknown and are currently under investigation.

## Figure legends

**Fig. S1. Ethanol-resistant cells grow in broth media**. Representative images of serial spot dilution of ethanol resistant sporulation assay of R20291_CM196_ and R20291_CM210_. Cells were spotted at 10-fold serial dilutions on BHIS agar plates supplemented with 0.1% sodium taurocholate. U: untreated cells, T: ethanol-treated cells.

**Fig. S2. Ethanol-resistant cells grow in agar media**. Representative images of serial spot dilution of ethanol resistant sporulation assay of R20291_CM196_ and R20291_CM210_. Cells were spotted at 10-fold serial dilutions on BHIS agar plates supplemented with 0.1% sodium taurocholate. U: untreated cells, T: ethanol-treated cells.

