## Supplementary material for "Characterization of a hypersporulating strain derivative of *Clostridioides difficile* R20291": Table S1

**Table S1.** Strains used in this work

| Strain | Description | Reference |
|---|---|---|
| $R20291_{CM210}$ | R20291 derivative strain | This work |
| $R20291_{CM196}$ | Hypersporulating R20291 derivative strain | This work |
