## Supplementary figures and images for "Characterization of a hypersporulating strain derivative of *Clostridioides difficile* R20291"

### Figure S1

#### Supplementary figure 1

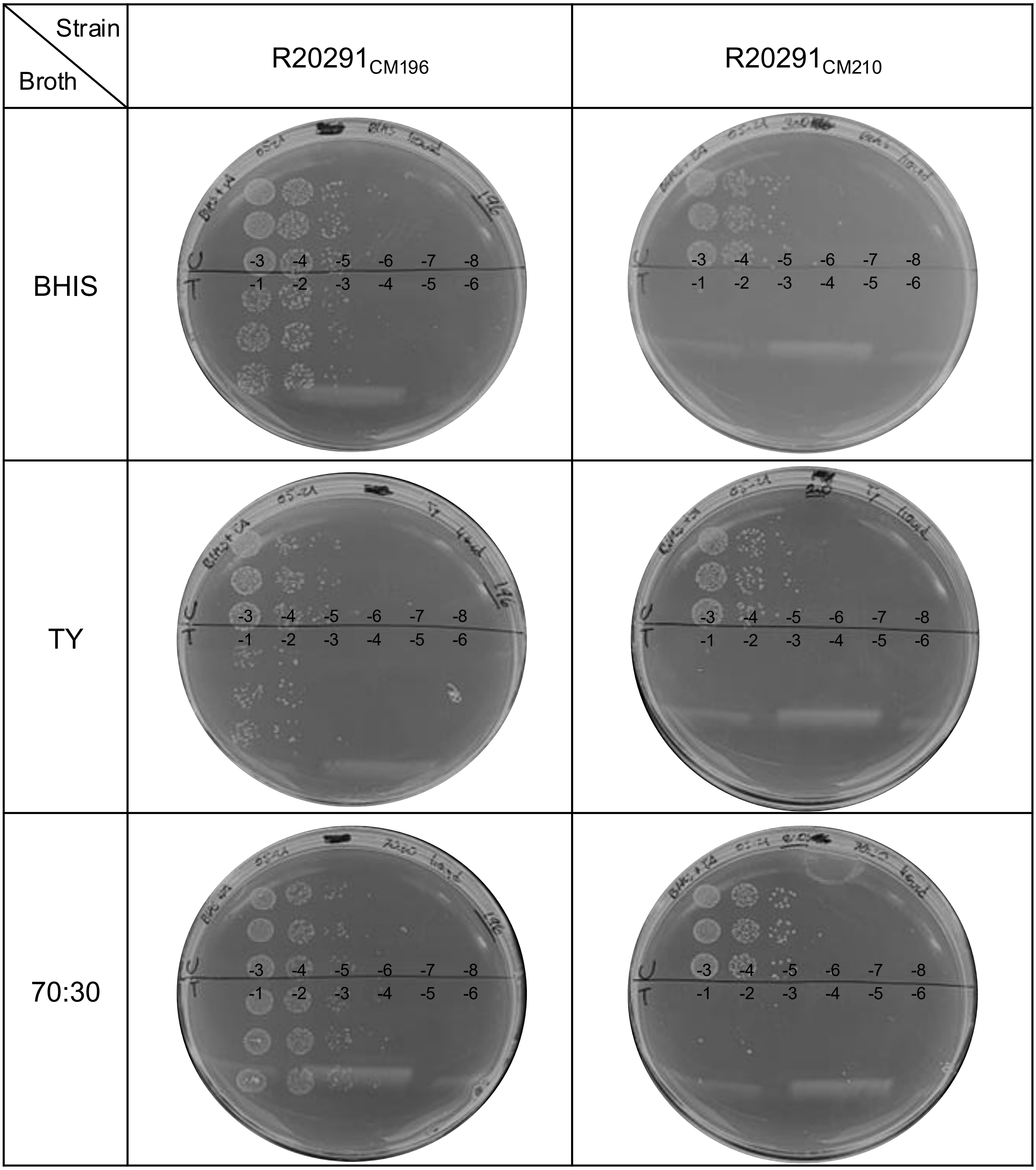

**Supplementary figure 2**

| Strain / Agar | $R20291_{CM196}$ | $R20291_{CM210}$ |
|---|---|---|
| BHIS | 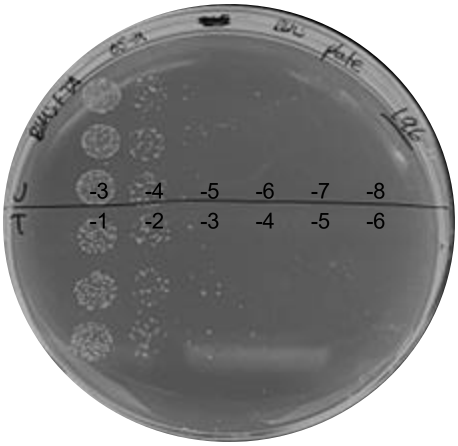 | 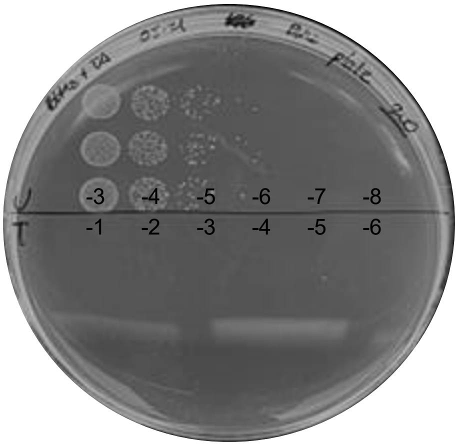 |
| TY | 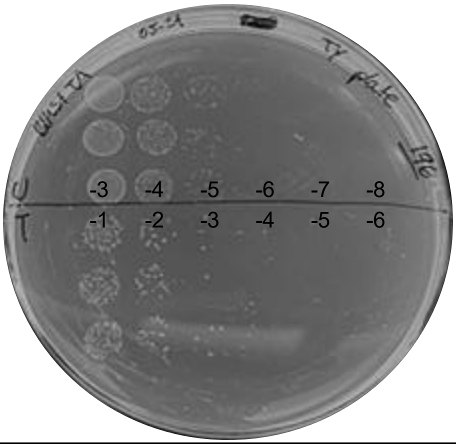 | 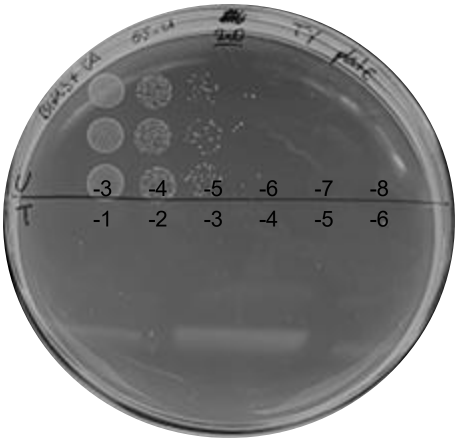 |
| 70:30 | 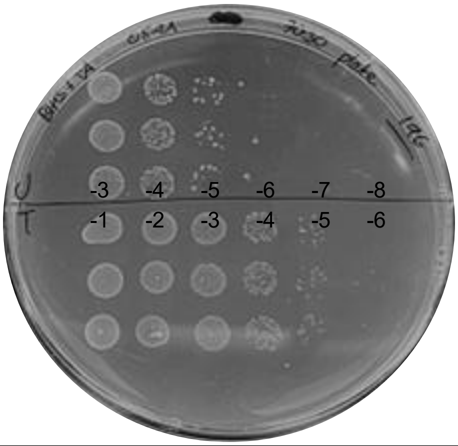 | 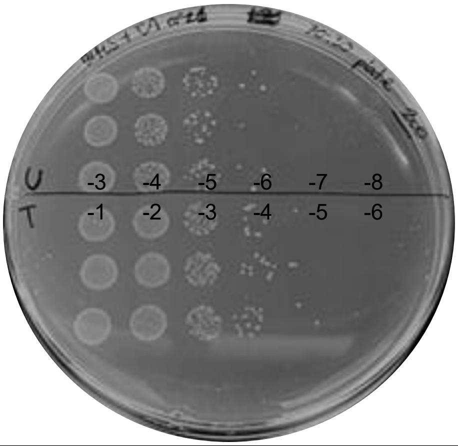 |
